# Gene editing of putative cAMP and Ca^2+^-regulated proteins using an efficient cloning-free CRISPR/Cas9 system in *Trypanosoma cruzi*

**DOI:** 10.1101/2023.07.09.548290

**Authors:** Miguel A. Chiurillo, Milad Ahmed, César González, Aqsa Raja, Noelia Lander

## Abstract

*Trypanosoma cruzi*, the agent of Chagas disease, must adapt to a diversity of environmental conditions that it faces during its life cycle. The adaptation to these changes is mediated by signaling pathways that coordinate the cellular responses to the new environmental settings. Cyclic AMP (cAMP) and Calcium (Ca^2+^) signaling pathways regulate critical cellular processes in this parasite, such as differentiation, osmoregulation, host cell invasion and cell bioenergetics. Although the use of CRISPR/Cas9 technology prompted reverse genetics approaches for functional analysis in *T. cruzi*, it is still necessary to expand the toolbox for genome editing in this parasite, as for example to perform multigene analysis. Here we used an efficient T7RNAP/Cas9 strategy to tag and delete three genes predicted to be involved in cAMP and Ca^2+^ signaling pathways: a putative Ca^2+^/calmodulin-dependent protein kinase (*CAMK*), Flagellar Member 6 (*FLAM6*) and Cyclic nucleotide-binding domain/C2 domain-containing protein (*CC2CP*). We endogenously tagged these three genes and determined the subcellular localization of the tagged proteins. Furthermore, the strategy used to knockout these genes allow us to presume that *TcCC2CP* is an essential gene in *T. cruzi* epimastigotes. Our results will open new venues for future research on the role of these proteins in *T. cruzi*.

## INTRODUCTION

Chagas disease is one of the 20 neglected tropical diseases affecting about 1 billion people in the world. This vector-borne infectious disease is caused by the protozoan parasite *Trypanosoma cruzi* and affects 6 to 7 million people in 21 Latin American countries, where the disease is endemic. This infection is considered a leading cause of disability and premature death in the Americas, although most affected individuals remain undiagnosed and untreated (WHO 2020). There is no vaccine to prevent Chagas disease or satisfactory treatment for chronic patients. If untreated, the infection persists for a lifetime, causing chronic cardiomyopathy or dilatation of the gastrointestinal tract in one-third of the infected population, several years or even decades after the initial contagion (Bonney et al. 2019). Understanding *T. cruzi* biology is crucial to develop alternative strategies to diagnose and treat this silent disease.

Developmental transformations involve key physiological processes for parasite survival. *T. cruzi* has a complex life cycle involving four main developmental stages that colonize very specific niches within its hosts, transitioning between stages in response to microenvironmental changes (Lander et al. 2021, Melo et al. 2020). The signal transduction pathways driving differentiation in *T. cruzi* are still poorly understood. Among them, cAMP signaling has been linked to the development of infective forms within the triatomine bug, a process called metacyclogenesis (Hamedi et al. 2015, Rangel-Aldao et al. 1988, Gonzales-Perdomo et al. 1988). cAMP has also been reported to mediate response to osmotic stress in *T. cruzi* (Schoijet et al. 2011, Rohloff et al. 2004, King-Keller et al. 2010, Docampo et al. 2013). In mammalian cells, the basic components of this signaling pathway are well defined, and the expression of these proteins in different microdomains determines spatiotemporal regulation of cAMP signals (Musheshe et al. 2018, Wang et al. 2022). We have recently identified two putative cAMP signaling microdomains in *T. cruzi*: the flagellar distal domain (flagellar tip) and the contractile vacuole complex (CVC), structures involved in cell adhesion and osmoregulation, respectively (Chiurillo et al. 2023). However, just a few components of this pathway have been characterized in *T. cruzi* (Lander et al. 2021, Schoijet et al. 2019).

Calcium ion (Ca^2+^) is another important intracellular signal in trypanosomatids that regulates essential cellular processes like host cell invasion (Moreno et al. 1994), differentiation (Lammel et al. 1996), osmoregulation (Rohloff et al. 2003, Dave et al. 2021), flagellar function (Engman et al. 1989), life/death decisions (Huang et al. 2013) and cell bioenergetics (Chiurillo et al. 2020, Chiurillo et al. 2017, Huang et al. 2013). As in vertebrate cells, cytosolic free Ca^2+^ concentration is strictly maintained in the range of 20-100 nm in different trypanosomatids (Moreno and Docampo 2003). However, most types of plasma membrane Ca^2+^ channel orthologs are absent in these protozoans (Docampo and Huang 2015). In the cytosol, soluble calcium-binding proteins sequester Ca^2+^ within different intracellular compartments. Some of them have been identified in the *T. cruzi* genome, but their role is still unknown.

The adaptation of the CRISPR/Cas9 system for genome editing to *T. cruzi*, has been a game change in the genetic manipulation of this parasite (Lander and Chiurillo 2019, Chiurillo and Lander 2021). However, the molecular methods available in *T. cruzi* still show limitations compared to those used in other pathogenic protozoa such as *Toxoplasma gondii* or *Trypanosoma brucei*. The lack of efficient inducible methods for loss-of-function analyses in *T. cruzi* restrains the investigation of the role of essential genes in this parasite. The absence of the RNAi machinery in *T. cruzi* (DaRocha et al. 2004) and the apparent difficulty in adapting methods for inducible protein depletion in this parasite, such as auxin-degron system, explain in part the mentioned limitations. Moreover, to the best of our knowledge, no large-scale or simultaneous multigene editing analysis have been reported in *T. cruzi.* Also, the lack of non-homologous end joining (NHEJ) machinery in trypanosomatids, makes necessary to use donor DNAs to induce homology-directed repair (HDR) after Cas9 cleavage for efficient gene editing, which limits large-scale approaches. Cloning-free systems to generate sgRNAs and donor DNA cassettes for delivery into *T. cruzi* cell lines genetically engineered to constitutively express Cas9 and T7 RNA polymerase (T7RNAP) could enable systematic analysis of multiple genes, as successfully done in *Leishmania* species (Damianou et al. 2020, Baker et al. 2021, Beneke et al. 2019). Although the CRISPR/Cas9/T7RNAP-mediated method has been used in *T. cruzi* (Costa et al. 2018, Roson et al. 2022, Pavani et al. 2020, de Lima et al. 2019), an approach that can be easily scaled up for multigene editing in this parasite is still needed.

Taking advantage of the efficacy of CRISPR/Cas9-medited genome editing in *T. cruzi, in* this study we aimed at developing a fast and scalable strategy to identify essential genes when null mutants are non-viable. We used this strategy to generate knockout and tagged cell lines of three genes predicted to be involved in Ca^2+^ and cAMP signaling pathways: *TcCAMK*, *TcFLAM6* and *TcCC2CP*. In addition, we performed a preliminary phenotype analysis of these mutants in *T. cruzi* epimastigotes.

## MATERIAL AND METHODS

### Chemicals and reagents

Alexa-conjugated secondary antibodies, HRP-conjugated secondary antibodies, Pierce ECL Western blotting substrate and BCA Protein Assay Kit were from Thermo Fisher Scientific Inc. Blasticidin S HCl was from Gibco. Puromycin was from Acros Organics (Fair Lawn, NJ). Anti-c-Myc (9E10) epitope tag monoclonal antibody was from Invitrogen. Benzonase® nuclease was from Novagen (EMD Millipore, Billerica, MA). GoTaq G2 Flexi DNA Polymerase, pGEM^®^-T Easy Vector Systems and T4 DNA Ligase were from Promega (Madison, WI). Restriction enzymes and Q5^®^ High-Fidelity DNA Polymerase were from New England Biolabs (Ipswich, MA). Fluoromount-G^®^ was from SouthernBiotech (Birmingham, AL). Nitrocellulose, polyacrylamide, and Precision Plus Protein Dual Color protein standard were from Bio-Rad (Hercules, CA). DNA oligonucleotides were purchased from Integrated DNA Technologies, Inc. (Coraville, Iowa). *E. coli* DH5α Mix & Go competent cells, ZymoPURE Plasmid Midiprep kit, and ZymoPURE Plasmid Miniprep kit were from Zymo Research (Irvine, CA). G418 Sulfate was from KSE Scientific (Durham, NC). Anti-tubulin mAb, mammalian cell protease inhibitor mixture (Sigma P8340), other protease inhibitors, and all other reagents of analytical grade were from Sigma (St. Louis, MO). The pMOTag23M vector (Oberholzer et al. 2006) was from Dr. Thomas Seebeck (University of Bern, Bern, Switzerland). Rabbit antibody against *T. brucei* vacuolar H-pyrophosphatase (TbVP1) (Lemercier et al. 2002) was from Dr. Norbert Bakalara (Ecole Nationale Supérieure de Chimie de Montpellier, Montpellier, France).

### Cell culture

*T. cruzi* Y strain epimastigotes were cultured in liver infusion tryptose (LIT) medium containing 10% heat-inactivated fetal bovine serum (FBS) at 28°C (Bone and Steinert 1956) T7/Cas9 cell line was maintained in medium containing 250 μg/ml G418. Mutant cell lines with single deleted allele were maintained in medium containing 250 μg/ml G418 and 5 μg/ml puromycin, while for KO cell lines 10 μg/ml blasticidin was additionally added. Cell density was determined using a Guava^®^ Muse^®^ Cell Analyzer (Luminex Corporation, Austin, TX).

### In silico analysis

Nucleotide sequences of *T. cruzi* Y strain *TcAEK1*(TcYC6_0120630), *TcCAMK* (TcYC6_0047690), *TcFLAM6* (TcYC6_0109380) and *TcCC2CP* (TcYC6_0079900) genes were retrieved from TriTrypDB genome data (tritrypdb.org). Prediction of protein domains of TcCAMK, TcFLAM6 and TcCC2CP amino acid sequences was made using web servers: https://ebi.ac.uk/interpro/ and https://prosite.expasy.org/scanprosite/. Selection of protospacers for knockout and tagging strategies was performed using EuPaGDT (eukaryotic pathogen CRISPR guide RNA/DNA design tool; http://grna.ctegd.uga.edu) (Peng and Tarleton 2015).

### Generation of a *T. cruzi* cell line stably expressing T7 RNA Polymerase and Cas9

To generate a *T. cruzi* cell line that constitutively expresses both T7RNAP and Cas9 nuclease (T7RNAP/Cas9), *T. cruzi* Y strain epimastigotes were transfected with circular pTREXn-T7RNAP/Cas9 plasmid, which was created by cloning the T7RNAP gene into pTREXn/Cas9 vector (Lander et al. 2015) upstream the Cas9-HA-GFP ORF, and interspaced by the *T. cruzi* trans-splicing region *HX1* (Fig. S1A). Clonal populations of parasites were obtained in 96-well pates by limiting dilutions in conditioned medium.

### Design and PCR-amplification of sgRNA templates

The 124-bp DNA templates for sgRNA transcription were designed as previously reported (Bassett and Liu 2014). The PCR contains a forward primer which is target-specific and includes the T7 polymerase binding site, the 20-nt of the target sequence or protospacer, and a region complementary to a common reverse primer (RvG00) containing the remaining sequence of the sgRNA (Table S1). sgRNA templates were amplified using Q5^®^ High-Fidelity DNA polymerase and 10 μM of each sgRNA-forward and RvG00 primers under the following conditions: a first denaturation step of 98°C for 30 sec, followed by 15 cycles of 98°C for 10 s, 55°C for 20 s, and 72°C for 15 s, then 25 cycles of 98 °C for 10 s, and 72°C for 20 s followed by a final extension 72°C for 2 min.

### Endogenous C-terminal gene tagging

The choice of the target sequence to design sgRNA templates to perform CRISPR/Cas9-mediated endogenous C-terminal tagging was done as previously described (Lander et al. 2017, Lander et al. 2016b). A donor DNA cassette containing the 3xc-Myc tag sequence and the puromycin resistance gene to induce HDR was amplified using the pMOTag23M vector (Oberholzer et al. 2006) as template. Primers to amplify by PCR the donor DNA cassette were designed to be 60-nt long, including the common nucleotide sequences to anneal to the pMOTag23M plasmid, and 39-nt and 34-nt target-specific sequences in the forward and reverse primers (Fw/Rv-CTag), respectively (Table S1). *T. cruzi* T7RNAP/Cas9 epimastigotes were co-transfected with the sgRNA template and the donor DNA, and then cultured for 2 weeks with G418 and puromycin for selection of resistant parasites. Endogenous gene tagging was verified by PCR from gDNA using primers Fw/Rv-CTag check (Table S1) and by western blot analysis using total protein extracts.

### CRISPR-Cas9 mediated gene deletion

Plasmids used as DNA template in PCR reactions to generate donor DNAs for gene deletion/disruption strategies were obtained by amplifying blasticidin S deaminase (*BSD*) and puromycin-N-acetyltransferase (*PAC*) genes and cloning them individually into pGEM®-T Easy Vector (Promega). The reverse primers to amplify both genes included GTGA as four last nucleotides, therefore GTGA is the 3’ end of *BSD* and *PAC* genes. The generated plasmids were named p*BSD* and p*PAC*, respectively. For PCR-amplification of resistance cassettes we used 60-nt primers that were designed as follows (Fw/Rv-KO, Table S1): The forward primers contain a 40-nt 5’UTR-containing homologous region (HR) plus 20 nucleotides of the plasmid backbone including the start codon (5’-GCCGCGGGAATTCGATTATG-3’). The reverse primers consist of 37 nucleotides of the 3’UTR-containing HR followed by 23 nucleotides of the plasmid backbone and the four last nucleotides of the antibiotic resistance genes, including the stop codon (5’-CGCGAATTCACTAGTGATTTCAC-3’). Homology donor targeting sequences in forward/reverse primers were designed by manual verification of the appropriate sequence in TriTrypDB. To amplify the donor DNA cassettes, 30 ng circular p*BSD* and p*PAC* plasmid DNA, 0.4 μM of 60-nt gene-specific forward and reverse primers, and 5% (v/v) DMSO when amplifying *PAC* cassette, and 1.5 U of GoTaq G2 Flexi DNA Polymerase, were mixed in 50 μl total volume. PCR steps were 2 min at 95°C, followed by 15 cycles of 20 s at 95°C, 20 s at 60°C, 55 s at 72°C, then other 25 cycles of 20 s at 95°C, 1 min at 72°C, followed by a final elongation step of 5 min at 72°C. Sizes of donor DNA cassettes are 515 bp and 715 bp for *BSD* and *PAC* resistant markers, respectively.

To generate single-KO or null mutant cell lines for each gene of interest we co-transfected *T. cruzi* T7RNAP/Cas9 epimastigotes with the sgRNA template and one (puromycin) or two (puromycin and blasticidin) donor DNAs, respectively. Transfected parasites were cultured for 2-3 weeks in the presence of G418 and puromycin, or G418, puromycin and blasticidin, for selection of single or double KO parasites, respectively. When no viable double KO cells were obtained by transfecting with two donor DNAs simultaneously, stable confirmed single-KO cells were subjected to a second round of transfection including the sgRNA template and the blasticidin resistance cassette. Gene disruption was verified using gDNA from mutant parasites by PCR using primers Fw/Rv-KO check (Table S1).

### Cell transfections

*T. cruzi* Y strain epimastigotes were transfected as described previously (Chiurillo et al. 2017). Briefly, *T. cruzi* epimastigotes in early exponential phase (4 x 10^7^ cells) were washed with PBS, pH 7.4, at room temperature (RT) and transfected in ice-cold CytoMix (120 mM KCl, 0.15 mM CaCl_2_, 10 mM K_2_HPO_4_, 25 mM HEPES, 2 mM EDTA, 5 mM MgCl_2_, pH 7.6) containing 25 μg of each plasmid construct and 25 μg of donor DNA in 4-mm electroporation cuvettes with three pulses (1500 volts, 25 microfarads) delivered by a Gene Pulser Xcell™ Electroporation System (Bio-Rad). Transfected epimastigotes were cultured in LIT medium supplemented with 20% heat-inactivated FBS until stable cell lines were obtained. When needed, the antibiotic (concentration) used for drug selection and maintenance was 250 μg/ml G418, 10 μg/ml blasticidin or 5 μg/ml puromycin.

### Western blot analysis

Western blots were carried as described previously (Lander et al. 2010). Briefly, parasites were harvested and washed twice in PBS and subsequently resuspended in radio-immunoprecipitation assay buffer (RIPA: 150 mM NaCl, 20 mM Tris-HCl, pH 7.5, 1 mM EDTA, 1% SDS, 0.1% Triton X-100) plus a mammalian cell protease inhibitor mixture (diluted 1:250), 1 mM phenylmethylsulfonyl fluoride, 2.5 mM tosyl phenylalanyl chloromethyl ketone (TPCK), 100 M *N*-(*trans*-epoxysuccinyl)-L-leucine 4-guanidinobutylamide (E64), and Benzonase Nuclease (25 units/ml of culture). The cells were then incubated for 1 h on ice and protein concentration was determined by BCA protein assay. Thirty micrograms of protein from each cell lysate were mixed with 6 Laemmli sample buffer (125 mM Tris-HCl, pH 7, 10% (w/v) ϕ3-mercaptoethanol, 20% (v/v) glycerol, 4.0% (w/v) SDS, 4.0% (w/v) bromphenol blue) before application to 10% SDS-polyacrylamide gels. Electrophoresed proteins were transferred onto nitrocellulose membranes with a Bio-Rad Trans-Blot^®^ Turbo™ Transfer System. Membranes were blocked with 5% nonfat dried skim milk in PBS-T (PBS containing 0.1% v/v Tween 20) overnight at 4 °C. Next, blots were incubated for 1 h, at RT, with a primary antibody (monoclonal anti-c-Myc-tag [1:1000] and monoclonal anti-tubulin [1:40,000]). After three washes with PBS-T, blots were incubated with the secondary antibody (goat anti-mouse IgG HRP-conjugated antibody, diluted [1:10,000]). Membranes were washed three times with PBS-T, then blots were incubated with Pierce ECL Plus Substrate and images were obtained and processed with a ChemiDoc Imaging System (Bio-Rad).

### Immunofluorescence analysis

Cells were washed with PBS and fixed with 4% paraformaldehyde in PBS for 1 h at RT. Cells were allowed to adhere to poly-L-lysine-coated coverslips and then permeabilized for 5 min with 0.1% Triton X-100. Permeabilized cells were blocked with PBS containing 3% BSA, 1% fish gelatin, 50 mM NH_4_Cl, and 5% goat serum overnight at 4 °C. Then, cells were incubated with a primary antibody (monoclonal anti-c-Myc-tag [1:100]), diluted in 1% BSA in PBS (pH 8.0) for 1 h at RT. Rabbit anti-TbVP1 antibodies were used at a dilution of 1:500. Cells were washed three times with 1% BSA in PBS (pH 8.0), and then incubated for 1 h at RT in the dark with Alexa Fluor 488 or Alexa Fluor 546-conjugated goat anti-mouse secondary antibodies (1:1000). Then, cells were washed and mounted on slides using Fluoromount-G mounting medium containing 5 μg/ml of 4,6-diamidino-2-phenylindole (DAPI) to stain DNA. Controls were done as described above but in the absence of a primary antibody. Differential interference contrast and fluorescence optical images were captured with a 100× objective in an Upright Motorized Fluorescence Microscope ECLIPSE Ni-E (Nikon) with NIS-Elements Advanced Research Package for 6D Image Acquisition and Analysis, and 2D fast deconvolution software.

## RESULTS

To generate a *T. cruzi* lineage that constitutively expresses Cas9 and T7RNAP we transfected Y strain epimastigotes with the pTREXn-T7RNAP/Cas9 construct (Fig S1A). pTREX vector integrates into the *T. cruzi* genome (Vazquez and Levin 1999), allowing the stable expression of T7RNAP and Cas9 genes. Expression of Cas9 in a clonal population was confirmed by western blot analysis and by direct observation of *T. cruzi* cells with green nuclei under fluorescence microscopy. T7RNAP/Cas9 epimastigotes showed no difference in growth compared to control (pTREXn empty vector) parasites (Fig S1B).

The cloning-free strategy for genome editing the T7RNAP/Cas9 system involves co-transfection of a sgRNA (with an upstream T7RNAP promoter) and donor DNA(s) as PCR products. In addition, to generate a null mutant population using the T7RNAP/Cas9 system in a single round of electroporation, it is necessary to transfect cells with 2 donor DNAs, each one containing a different resistance marker. To streamline strategy, we generated a set of 2 plasmids using the pGEM^®^-T Easy Vector Systems (Promega) as backbone to clone blasticidin and puromycin resistance genes (p*BSD* and p*PAC*, respectively). The design allows to use a single primer set to amplify any of the resistance cassettes. Primers to amplify by PCR the donor DNA cassettes were designed to be 60-nt long, including the common nucleotide sequences to anneal on p*BSD* and p*PAC* plasmids, and 40-nt or 37-nt target-specific HR sequence in the forward and reverse primers (Fw/Rv-KO, Table S1), respectively. Primers with 30-nt HRs were also tested showing similar efficiency. Our strategy is based on choosing the HR in the 5’ and 3’ UTRs immediately flanking the start and stop codon of the gene of interest (GOI), respectively (Fig. 1A). In the present study we followed a two-step strategy for gene deletion: 1) in a first round of transfections, we transfected in duplicates T7RNAP/Cas9 epimastigotes with an specific sgRNA template and one (*PAC*) or two (*PAC* and *BSD*) donor DNAs (Fig. 1A); 2) if the KO cell line was not viable, we performed a second round of transfection to deliver the sgRNA template plus the *BSD* cassette in a stable single allele deletion mutant cell line (single KO) which was generated in the first round of transfections.

**FIGURE 1.**
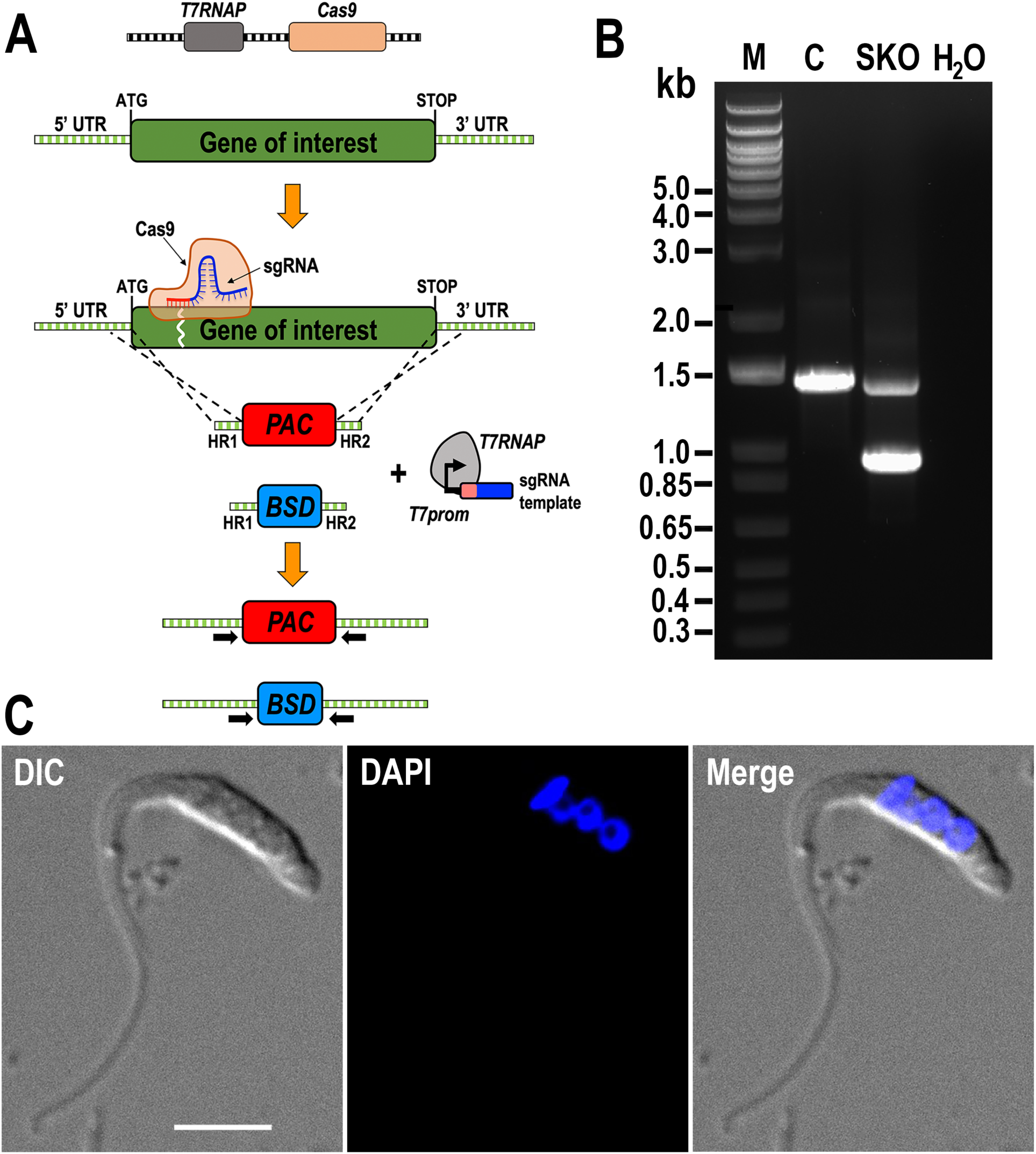
T7RNAP/Cas9 system for genome editing in *Trypanosoma cruzi*. A) Schematic representation of the general strategy used for CRISPR/Cas9-mediated gene deletion in *T. cruzi* cell line expressing T7RNAP and Cas9. This system requires constitutive expression of Cas9 and T7RNAP; while sgRNA and donor DNAs are PCR-generated. The scheme shows the integration of the repair cassettes containing homologous regions (HR1 and HR2) after the double strand break generated by the ribonucleoprotein. sgRNA template including a T7 promoter is transfected along with donor DNAs. UTR, untranslated region; *PAC*, puromycin N-acetyltransferase; *BSD*, Blasticidin S deaminase. Horizontal black arrows indicate primers used for checking integration of donor DNA by PCR. B) Only one *TcAEK1* allele was disrupted at its genomic locus in the SKO cell lines. Lanes: M, 1 Kb Plus ladder; C, T7RNAP/Cas9 control; SKO, *TcAEK1*-*SKO*; H_2_O, PCR negative control. PCR fragment sizes: WT *TcAEK1*, 1,436 bp; *PAC*-replaced allele, 968 bp. C) Representative images displaying aberrant cell morphology and DNA content of a *TcAEK1*-*SKO* epimastigote. DIC, differential interference contrast. Merge image shows DAPI staining (blue) and DIC. Scale bar: 5 μm.

To validate the effectiveness of our T7RNAP/Cas9 system we attempted to generate a null mutant cell line of the AGC essential kinase 1 (*TcAEK1*; TriTrypDB ID: TcYC6_0120630; 1179 bp). *AEK1* in *T. cruzi* is essential for epimastigote proliferation, trypomastigote host cell invasion, and amastigote replication, as well as in cytokinesis regulation (Chiurillo et al. 2021). Using a previous CRISPR/Cas9 method we adapted for *T. cruzi* (Lander et al. 2015, Lander et al. 2016a, Chiurillo et al. 2020, Chiurillo et al. 2019, Chiurillo et al. 2017, Negreiros et al. 2021, Lander and Chiurillo 2019), which involves the constitutive expression of Cas9 and the sgRNA, it was only possible to generate a single *TcAEK1* allele deletion cell line. In this previous work, several attempts with different strategies demonstrated the essentiality of *TcAEK1* because it was not possible to obtain null mutants. Consistently, here we were just able to obtain a *TcAEK1* single KO (*TcAEK1*-*SKO*) cell line (Fig. 1B), since both attempts, transfecting with 2 resistance cassettes at the same time or in two sequential rounds of transfections, were unsuccessful in generating null mutant cells for this gene. As expected, *TcAEK1*-*SKO* epimastigotes showed a defect in cytokinesis (Fig. 1C).

We next investigated a putative Ca^2+^/calmodulin-dependent protein kinase (*TcCAMK*; TriTrypDB ID: TcYC6_0047690; 1638 bp), whose ortholog in *T. brucei* is an essential gene involved in cytokinesis (Jones et al. 2014). Initially we evaluated the T7RNAP/Cas9 system for endogenous C-terminal tagging method by inserting the c-Myc epitope tag sequence into the 3’ end of *TcCAMK* gene. The strategy consisted in co-transfecting T7RNAP/Cas9 epimastigotes with a specific sgRNA template targeting the 3’ end of *TcCAMK* gene and a DNA donor molecule for each gene amplified from the pMOTag23M vector (Lander et al. 2017, Oberholzer et al. 2006). In general, this strategy will tag one of the alleles of the gene (Fig. 2A). Using this method *TcCAMK* was efficiently tagged at the endogenous locus with a 3xc-Myc tag, as confirmed by PCR (Fig. 2B), and western blot analysis using anti-c-Myc antibodies, detecting a band of the predicted size (67 kDa) (Fig. 2C). Immunofluorescence analysis (IFA) of the mutants showed that TcCAMK has a cytosolic localization in *T. cruzi* epimastigotes (Fig. 2D). Subsequently, we generated *TcCAMK*-*KO* epimastigotes by transfecting *T. cruzi* epimastigotes simultaneously with *BSD* and *PAC* resistance cassettes. *TcCAMK*-*KO* epimastigotes, confirmed by PCR (Fig. 2E), showed no difference in growth rate compared to control parasites (Fig. 2F), indicating that this putative *TcCAMK* gene is not essential at least for the epimastigote stage of *T. cruzi*.

**FIGURE 2.**
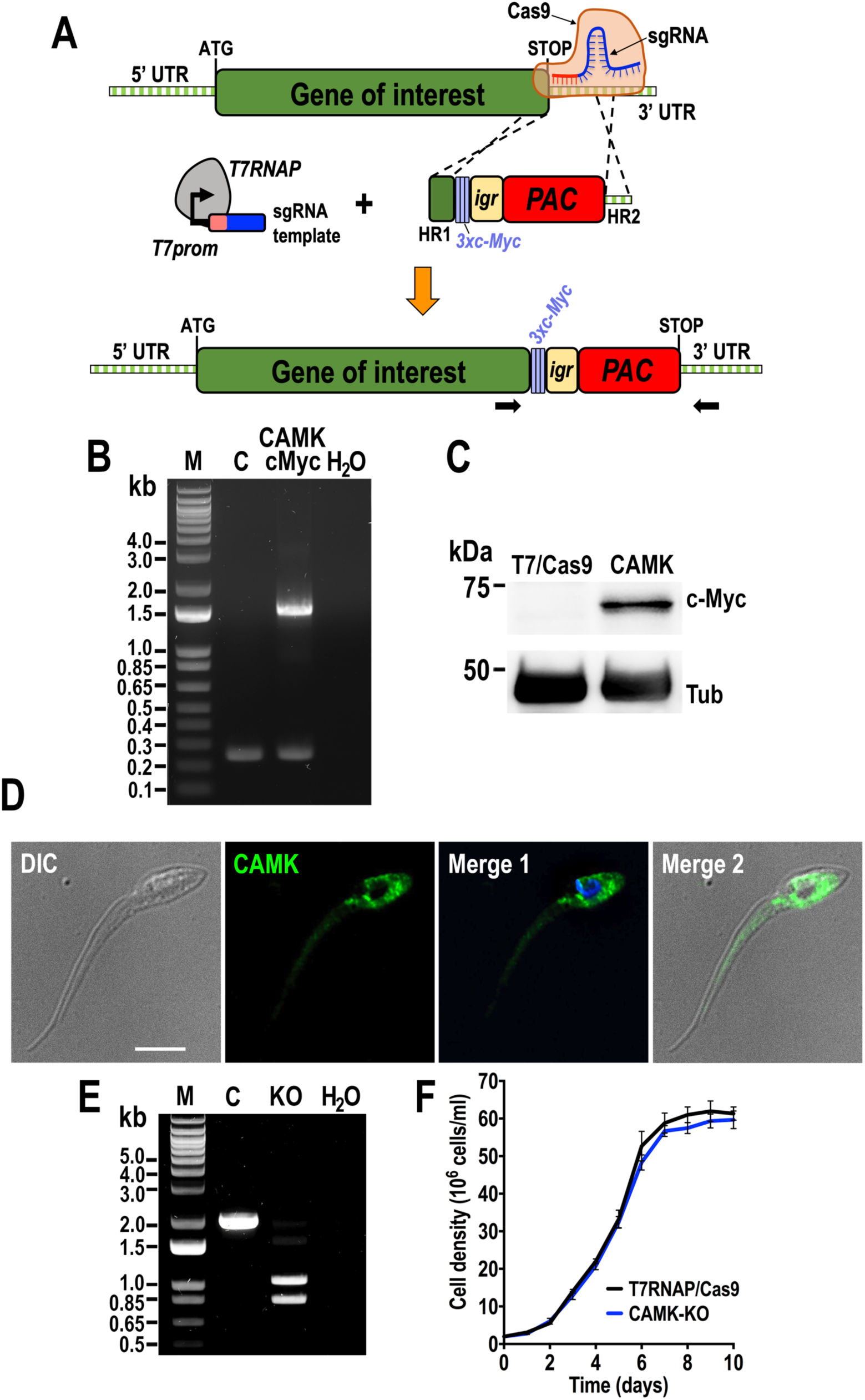
*TcCAMK* genome editing using a T7RNAP/Cas9 system. A) Schematic representation of the general strategy used for CRISPR/Cas9-mediated C-terminal endogenous tagging. *T. cruzi* epimastigotes constitutively expressing T7RNAP and Cas9 are transfected with two DNA fragments obtained by PCR: sgRNA template including a T7 promoter designed to target the 3’ end of the gene of interest, and a donor DNA including homologous regions (HR1 and HR2) flanking a puromycin resistance cassette, to induce homologous direct repair. UTR, untranslated region; *Igr*, Tubulin intergenic region; *PAC*, puromycin N-acetyltransferase. Horizontal black arrows indicate primers used for checking integration of donor DNA by PCR. B) PCR verification of integration of donor DNA in the 3’ end of the endogenous *TcCAMK* locus. *TcCAMK* allele sizes: WT, 240 bp; 3xc-Myc-tagged, 1,520 bp. Lanes: M, 1 Kb Plus DNA ladder; C, T7RNAP/Cas9 control; CAMK-cMyc, *TcCAMK*-3xc-Myc; H_2_O, PCR negative control. C) Western blot of TcCAMK-3xc-Myc (CAMK) using antibodies anti-c-Myc. Protein extract of the parental T7RNAP/Cas9 strain (T7/Cas9) was used as control. Tubulin was used as a loading control. D) Localization of endogenously tagged TcCAMK-3xc-Myc in epimastigotes using anti-c-Myc antibodies. DIC, differential interference contrast. CAMK (green), *TcCAMK*-3xc-Myc. Merge 1 image shows *TcCAMK*-3xc-Myc (green) and DAPI staining (blue), and Merge 2 shows *TcCAMK*-3xc-Myc (green) and DIC. Scale bar: 5 μm. E) PCR verification of resistance cassette integration in both *TcCAMK* alleles. Lanes: M, 1-kb Plus DNA ladder; C, T7RNAP/Cas9 control; KO, *TcCAMK*-KO; H_2_O, PCR negative control. PCR fragment sizes: WT *TcCAMK*, 1,960 bp; *PAC*-replaced allele, 1050 bp, *BSD* (Blasticidin S deaminase)-replaced allele, 850 bp. F) Growth of control (T7RNAP/Cas9) and *TcCAMK*-KO (CAMK-KO) epimastigotes in LIT medium (*n* = 3).

Using similar strategies, we studied the subcellular localization of TcFLAM6 (Flagellar Member 6, TriTrypDB: TcYC6_0109380, 4191 bp), whose ortholog was identified as a flagellar protein of *T. brucei* procyclic forms (Subota et al. 2014) and it is predicted to have two cyclic nucleotide-monophosphate binding domains or CBD (Jager et al. 2014). Fig. 3 shows the efficient tagging of *TcFLAM6* with 3xc-Myc at the endogenous locus, as detected by PCR, western blot analysis (predicted 162 kDa band) and IFA (Fig. 3A-C). Interestingly, TcFLAM6 localized to the proximal region of the epimastigote flagellum (Fig. 3C). *TcFLAM6*-*KO* cell line was generated after transfecting stable puromycin-resistant *TcFLAM6*-*SKO* epimastigotes with the sgRNA template and *BSD* donor DNA, since no viable *TcFLAM6* null mutants were obtained when the two donor resistance cassettes were delivered at the same time. Replacement of both alleles in *TcFLAM6*-*KO* cells was confirmed by PCR (Fig. 3D). Moreover, *TcFLAM6*-*KO* epimastigotes showed a significant lower growth rate in LIT medium in the exponential phase of growth, compared to control cells (Fig. 3E).

**FIGURE 3.**
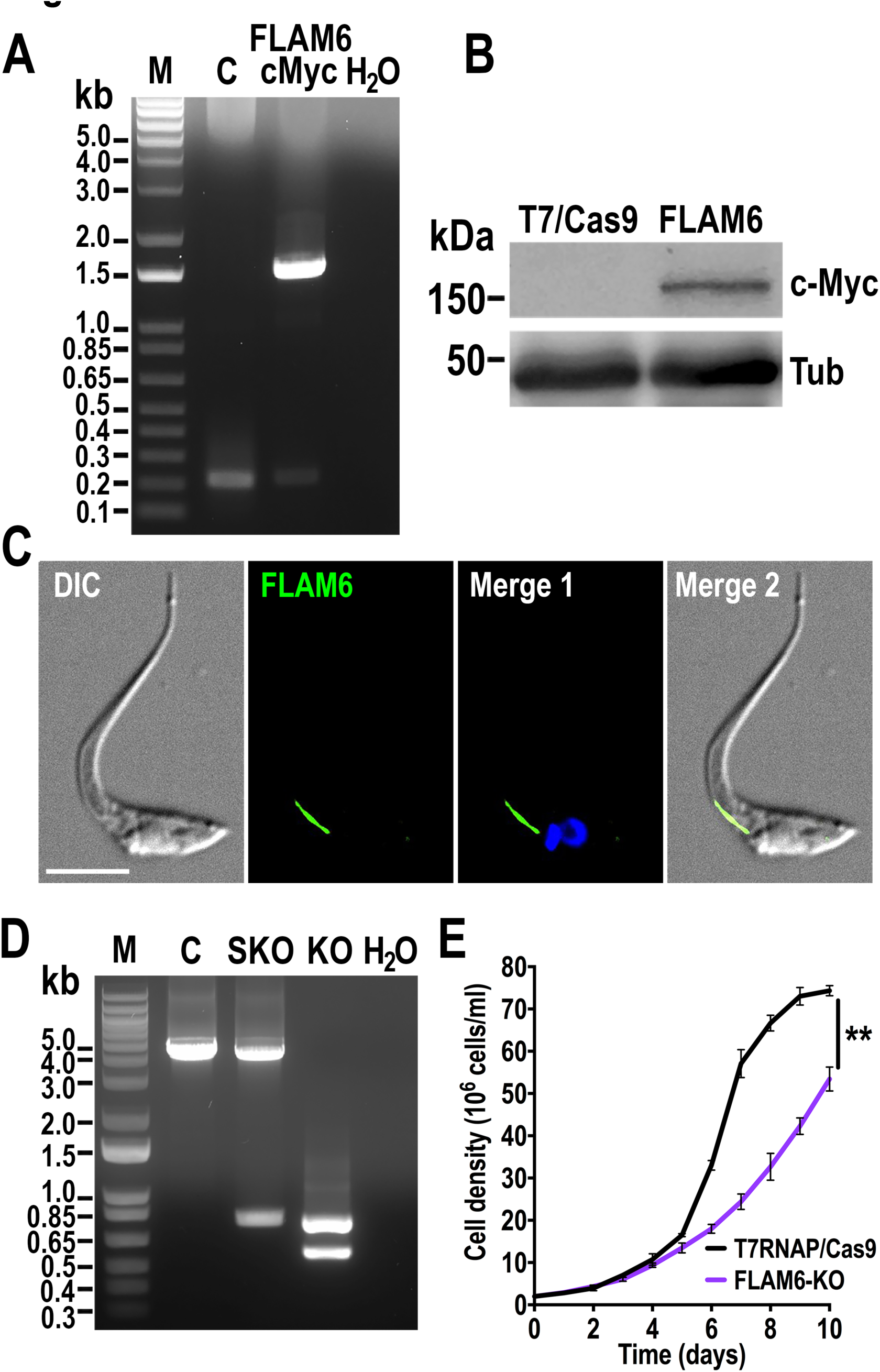
*TcFLAM6* genome editing using a T7RNAP/Cas9 system. A) PCR verification of integration of DNA donor at the 3’ end of the endogenous *TcFLAM6* locus. *TcFLAM6* allele sizes: WT, 215 bp; 3xc-Myc-tagged, 1,476 bp. Lanes: M, 1 Kb Plus DNA ladder; C, T7RNAP/Cas9 control; FLAM6-cMyc, *TcFLAM6*-3xc-Myc; H_2_O, PCR negative control. B) Western blot of TcFLAM6-3xc-Myc (FLAM6) using antibodies anti-c-Myc. Protein extract of the parental T7RNAP/Cas9 (T7/Cas9) cell line was used as control. Tubulin was used as a loading control. C) Localization of endogenously tagged TcFLAM6-3xc-Myc in epimastigotes using anti-c-Myc antibodies. DIC, differential interference contrast. FLAM6 (green), *TcFLAM6*-3xc-Myc. Merge 1 image shows *TcFLAM6*-3xc-Myc (green) and DAPI staining (blue), and Merge 2 shows *TcFLAM6*-3xc-Myc (green) and DIC. Scale bar: 5 μm. D) PCR verification of resistance cassette integration in *TcFLAM6* alleles. Lanes: M, 1-kb Plus DNA ladder; C, T7RNAP/Cas9 control; SKO, *TcFLAM6*-*SKO*; KO, *TcFLAM6*-*KO*; H_2_O, PCR negative control. PCR fragment sizes: WT *TcFLAM6*, 4,252 bp; *PAC* (puromycin N-acetyltransferase)-replaced allele, 776 bp, *BSD* (Blasticidin S deaminase)-replaced allele, 576 bp. E) Growth of control (T7RNAP/Cas9) and *TcFLAM6*-KO (FLAM6-KO) epimastigotes in LIT medium. Values represent means ± SD of three independent experiments (\*\**p* < 0.01, Student’s t-test).

Finally, we investigated a putative cAMP binding protein (TriTrypDB: TcYC6_0079900), which is 5076-bp gene and whose predicted protein product (∼187 kDa) contains two N-terminal CBDs and three C2 domains (calcium binding motifs) in the C-terminal region, according to InterPro (ebi.ac.uk/interpro/) (Fig. S2A). We named this protein as <u>C</u>yclic nucleotide-binding domain/<u>C2</u> domain-<u>C</u>ontaining <u>P</u>rotein (*TcCC2CP*). Then, we generated a *TcCC2CP* endogenous C-terminal tagged cell line, as confirmed by PCR and western blot analysis (Fig. 4A,B). *TcCC2CP*-3xc-Myc protein shows vesicle localization by IFA, with a pattern compatible with cytoplasmic organelles or endomembranes (Fig. 4C). Because the localization of TcCC2CP observed in epimastigotes resembles that exhibited by acidocalcisome-resident proteins (Lander et al. 2016b, Huang et al. 2014), and the predicted interaction of TcCC2CP with Ca^2+^, we performed an IFA using antibodies against the vacuolar proton pyrophosphatase (VP1), an acidocalcisome marker (Lemercier et al. 2002) and anti-c-Myc antibodies, to determine whether both proteins co-localize in this organelle. A partial co-localization was observed between TcCC2CP and TcVP1 in epimastigotes (Fig. S2B). However, additional analyses in other developmental stages, and using additional subcellular markers, should be performed to confirm this localization, such as markers of glycosomes or vesicular trafficking components. Furthermore, no populations of *TcCC2CP* null mutants were recovered after simultaneous transfection with two donor DNA cassettes or after performing a second round of transfection using *TcCC2CP*-*SKO* cell line, which could be easily generated (Fig. 4D). Interestingly, *TcCC2CP*-*SKO* epimastigotes exhibited a significantly slower growth in LIT medium than the parental T7RNAP/Cas9 cell line (Fig. 4E).

**FIGURE 4.**
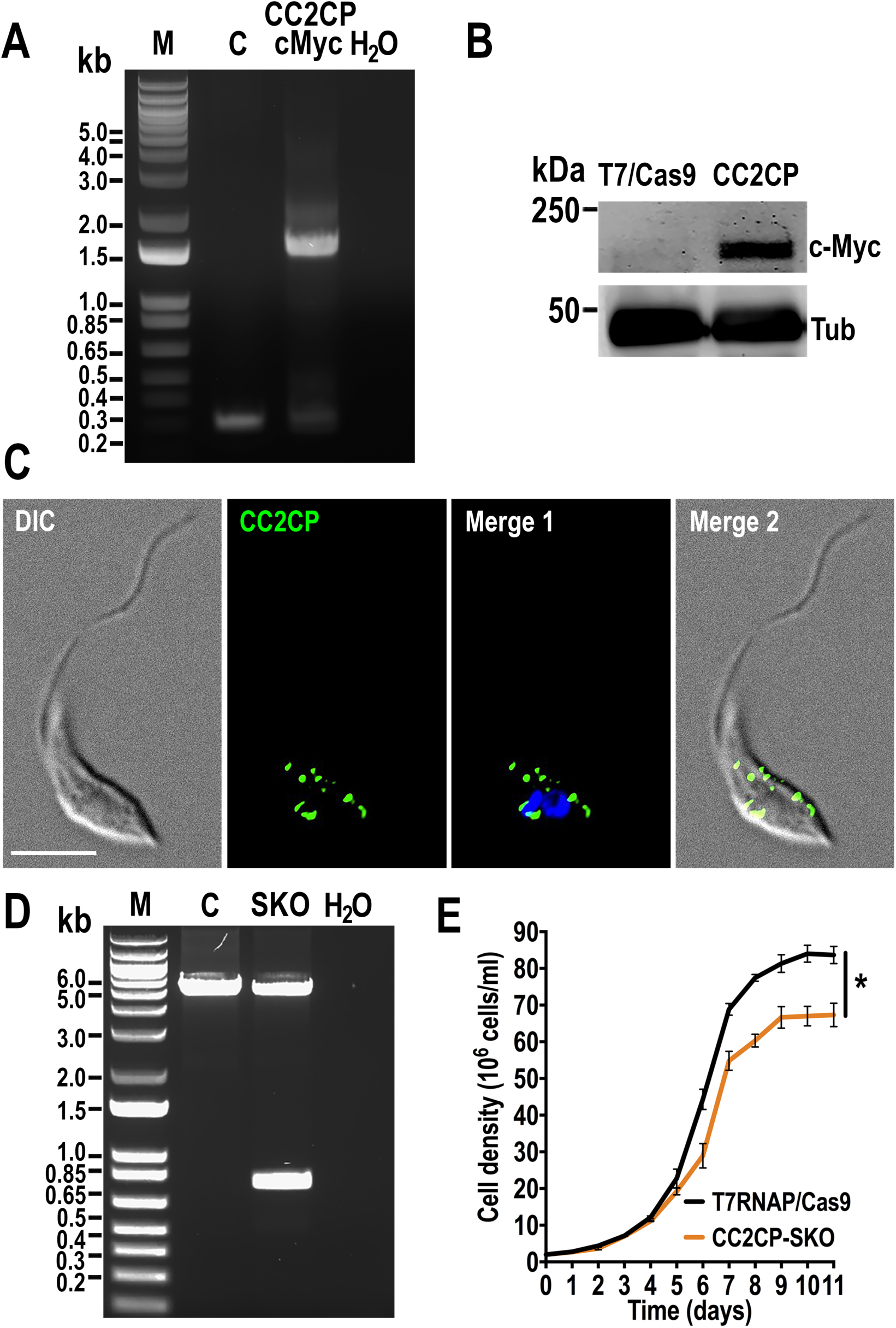
*TcCC2CP* genome editing using a T7RNAP/Cas9 system. A) PCR verification of integration of DNA donor in the 3’ end of the endogenous *TcCC2CP* locus. *TcCC2CP* allele sizes: WT, 298 bp; 3xc-Myc-tagged, 1,578 bp. Lanes: M, 1-kb Plus DNA ladder; C, T7RNAP/Cas9 control; CC2CP-cMyc, *TcCC2CP*-3xc-Myc; H_2_O, PCR negative control. B) Western blot of TcCC2CP-3xc-Myc (CC2CP) using antibodies anti-c-Myc. Protein extract of the parental T7RNAP/Cas9 cell line was used as control. Tubulin was used as a loading control. C) Localization of endogenously tagged TcCC2CP-3xc-Myc in epimastigotes using anti-c-Myc antibodies. DIC, differential interference contrast. CC2CP (green), *TcCC2CP*-3xc-Myc, Merge 1 image shows *TcCC2CP*-3xc-Myc (green) and DAPI staining (blue), and Merge 2 shows *TcCC2CP*-3xc-Myc (green) and DIC. Scale bar: 5 μm. D) Only one *TcCC2CP* allele was disrupted at its genomic locus in the SKO cell line. Lanes: M, 1 Kb Plus DNA ladder; C, T7RNAP/Cas9 control; SKO, *TcCC2CP*-SKO; H_2_O, PCR negative control. PCR fragment sizes: WT *TcCC2CP*, 5,210 bp; *PAC* (puromycin N-acetyltransferase)-replaced allele, 745 bp. E) Growth of control (T7RNAP/Cas9) and *TcCC2CP*-*SKO* (CC2CP-SKO) epimastigotes in LIT medium. Student’s t test was applied to compare growth rates calculated from each growth curve (*n* = 3; ***, *p* < 0.001).

## DISCUSSION

In *T. cruzi*, the CRISPR/Cas9 strategy involving co-transfection of epimastigotes with a vector for constitutive expression of Cas9 and sgRNA, plus a donor DNA cassette to induce HDR, has been successfully used to generate gene knockout mutants and to perform endogenous gene tagging (Lander et al. 2015, Lander et al. 2016b, Lander and Chiurillo 2019, Chiurillo et al. 2020, Chiurillo et al. 2019, Bertolini et al. 2019, Negreiros et al. 2021, Chiurillo et al. 2021). However, this strategy can exhibit some drawbacks, such as long selection time to obtain stable KO cell lines and the need of cloning the sgRNA into the Cas9/pTREXn vector before transfection. Moreover, the constitutive expression of Cas9 and sgRNA has been associated to off-target effects (Hsu et al. 2013, Hajiahmadi et al. 2019).

Other successful CRISPR/Cas9-mediated genome editing methods used in *T. cruzi* involve transfection of *in vitro* transcribed sgRNA(s), either with constitutive expression of Cas9 (Peng et al. 2014, Marek et al. 2021, Marcelino et al. 2023, Burle-Caldas et al. 2022) or delivered as an RNP complex with recombinant Cas9 (Soares Medeiros et al. 2017, Saenz-Garcia et al. 2021). In the latter cases, the transient presence of the sgRNA (as well as Cas9 when it is transfected as RNP complex) is considered beneficial. However, these strategies are not suitable for large-scale gene editing. CRISPR/Cas9 strategies involving transfection of epimastigotes stably expressing Cas9 and T7 RNA polymerase, which drives sgRNA transcription *in vivo* from DNA templates generated by PCR, plus donor cassette(s) for HDR, have significantly boosted reverse genetics studies in *Leishmania* (Jones et al. 2018, Grewal et al. 2019, Damianou et al. 2020, Turra et al. 2021, Martel et al. 2017, Espada et al. 2021, Beneke et al. 2017), and it was also used in *T. brucei* (Beneke et al. 2017). This cloning-free strategy has substantially increased the number of genes studied in different *Leishmania* species, mainly because the method can be used for simultaneously editing multiple genes in a large-scale format (Damianou et al. 2020, Baker et al. 2021, Beneke et al. 2019). Moreover, the transient expression of the sgRNA represents another advantage of the method to avoid off-target effects. This CRISPR/T7RNAP/Cas9-mediated genome editing strategy has been used in *T. cruzi* CL Brener strain for gene tagging, gene mutation and ablation of a few genes (Costa et al. 2018, Roson et al. 2022, Pavani et al. 2020, de Lima et al. 2019). The strategy followed to perform gene deletion in the present work shows some differences with the previous reported: 1) T7RNAP/Cas9 expressing cell line was generated by transfecting both genes cloned into the *T. cruzi-*specific expression vector pTREXn, instead of pLew13 plasmid; 2) epimastigotes are transfected with one sgRNA template instead of two; 3) Alleles of the GOI are replaced only by the resistance marker gene(s), preserving the endogenous UTRs to regulate their expression. Given the absence of the RNAi machinery in *T. cruzi*, a cloning-free CRISPR/T7RNAP/Cas9-mediated strategy seems to be the most appropriate approach currently available to perform large-scale gene editing in this parasite.

Furthermore, when a gene knockout attempt fails with any genome editing method, it often leads to the conclusion that the gene is “essential”. However, possible technical issues such as potential mismatches in the sequences of the protospacer and the protospacer adjacent motif (PAM) or incorrect assembly of the reference genome, can make the locus non-editable. Instead of repeating transfections or designing/selecting new sgRNA and homology regions (HRs), in this work per each GOI we transfected parasites with either one or two resistance cassettes, so that if double resistant parasites were not obtained, stable single KO cells were transfected with another DNA resistance marker in a second round of transfection. Using the described strategy, by obtaining at least single allele mutants, we can also ensure that the engineered sgRNA is efficient in targeting the GOI for Cas9-mediated cleavage. Our previous experience indicates that single KO cell lines can be generated even in cases in which mutant epimastigotes exhibit lower growth rates compared with parental cells (Lander et al. 2018, Dos Santos et al. 2021, Chiurillo et al. 2021). Following the current strategy as a routine for gene ablation we were able to: 1) obtain null mutants faster than performing multiple attempts to delete both alleles at the same time, and 2) be more confident in concluding that a gene is essential in *T. cruzi* epimastigotes. In addition, considering the limited number of effective resistance markers available for selection of *T. cruzi* mutants, endogenous genes could be restored in KO parasites, as the hygromycin resistance marker is still available for selection of add back mutants.

In this work we used a CRISPR/T7RNAP/Cas9 system to analyze three different genes, for which null mutant epimatigotes were obtained in the first attempt for *TcCAMK*, or after transfecting single null parasites for *TcFLAM6*. However, we failed to obtain *TcCC2CP*-*KO* cells even after performing a second round of transfection using the *TcCC2CP*-*SKO* cell line. Interesting, the results correlate with fact that *TcFLAM6*-*KO* and *TcCC2CP*-*SKO* epimastigotes exhibited lower growth rates than the parental cell line, which suggests both genes are important for *T. cruzi* in this developmental stage.

Ca^2+^/calmodulin-dependent protein kinases (CAMK) comprise a group of Serine/Threonine protein kinases that are regulated by allosteric activation with Ca^2+^/calmodulin when the intracellular concentration of Ca^2+^ increases (Swulius and Waxham 2008). The CAMK group (which includes the AMPK) is poorly represented in trypanosomatid genomes compared to the human kinome (Parsons et al. 2005). In the *T. cruzi* YC6 genome available in TriTrypDB we found 15 putative CAMK and CAMKL (CAMK-like) genes, among which we are particularly interested in one of them (TriTrypDB ID: TcYC6_0047690), whose ortholog in *T. brucei* (TriTrypDB ID: Tb927.7.6220) was previously assessed in a kinome-wide RNAi library (Jones et al. 2014). In that work, the knockdown of *TbCAMK* induced growth arrest and cytokinesis defects in bloodstream forms. However, different to *TbCAMK-*downregulated cells, *TcCAMK-KO* epimastigotes did not exhibit morphological alterations or a growth defect, suggesting that the *T. cruzi* ortholog is not essential at least in the epimastigote stage.

In a proteomic analysis of flagella of *T. brucei* procyclic forms were identified eight novel proteins, FLAM1 to 8 (Subota et al. 2014), which were recently also detected in an APEX-based proximity proteomics approach in the same stage of this parasite (Velez-Ramirez et al. 2021). Remarkably, FLAM6 contains two CBDs suggesting that this protein could bind cAMP and eventually have a role in the flagellar function as an effector of this second messenger in trypanosomatids. Here, we observed that TcFLAM6 is localized in the proximal third of the epimastigote’s flagellum, similar to what was observed in *T. brucei* procyclic forms in which TbFLAM6 is restricted to the proximal portion of the flagellum, independently of its length (Subota et al. 2014). The fact that in procyclic forms, TbFLAM6 is restricted to the proximal half of the axoneme at all stages of formation of the new flagellum, even when the flagellum starts to elongate, suggests a strict spatiotemporal regulation of FLAM6 synthesis and flagellar assembly (Subota et al. 2014). Moreover, *TcFLAM6*-*KO* epimastigotes exhibited a lower growth rate in the exponential phase compared to parental cells, which in addition to the unsuccessful generation of a null mutant in a first round of transfection with two resistance markers, confirms the importance of this gene in *T. cruzi* epimastigotes. Interestingly, in a proximity-dependent biotinylation approach of flagellar proteins of *T. cruzi*, unlike other FLAM proteins (FLAM1,2,3,5,7 and 8), TcFLAM6 was only detected in epimastigotes and not in intracellular amastigotes (Won et al. 2023).

TcCC2CP predicted CBDs were previously analyzed *in silico* in a bioinformatic identification of cyclic nucleotide binding proteins in *T. cruzi* (Jager et al. 2014). Interestingly this protein also has C2 domains, which are structural domains involved in targeting proteins to cell membranes in a Ca^2+^-dependent manner (Zhang and Aravind 2010). Furthermore, transcriptomic analyses revealed higher expression levels of *TcCC2CP* in metacyclic trypomastigotes in *T. cruzi* Brazil and Dm28c strains (Minning et al. 2009, Smircich et al. 2015). In this work *TcCC2CP* could not be ablated, and *TcCC2CP*-*SKO* parasites exhibited a growth defect in epimastigotes, suggesting that *TcCC2CP* is an essential gene in these insect forms, and could play an important role in other developmental stages of *T. cruzi.* Many proteins containing C2 domains are implicated in signal transduction or membrane trafficking (Nalefski and Falke 1996). The presence of both CBD and C2 domains and the subcellular localization observed by IFA suggest that TcCC2CP is a component of the cAMP signaling pathway coupled with molecules or organelle trafficking.

In summary, the gene editing strategies developed in this work expand the toolkit available to perform reverse genetics in *T. cruzi*. Particularly, the T7RNAP/Cas9 system generated, and the strategy used to obtain gene deletion mutants, could facilitate large-scale gene analyses in this parasite. Our preliminary study of two putative cAMP binding proteins, TcFLAM6 and TcCC2CP, the latter containing also predicted C2 domains, suggest the importance of them at least in the epimastigote stage of *T. cruzi*. Further work, evaluating the expression pattern, interactome and role of these proteins in other developmental stages are necessary to elucidate their importance in *T. cruzi* life cycle.

## ACKNOWLEDGMENTS

Funding for this work was provided by the National Institute of Allergy and Infectious Diseases of the National Institutes of Health (Award number R00AI137322 to N. Lander) and the Latino Faculty Association of University of Cincinnati (2023 LFA award to M.A. Chiurillo). A. Raja was an UPRISE awardee at the University of Cincinnati.

## SUPPORTING INFORMATION

**FIGURE S1**

T7RNAP/Cas9 system for genome editing in *T. cruzi*. A) pTREXn-T7RNAP/Cas9 vector map. B) Growth of empty vector (pTREXn) and T7RNAP/Cas9 *T. cruzi* epimastigotes in LIT medium (*n* = 3).

**FIGURE S2**

TcCC2CP topology and localization. A) Schematic representation of the TcCC2CP topology. The predicted amino acid sequence of *TcCC2CP* (TriTrypDB ID: TcYC6_0079900) was scanned by InterProScan (https://www.ebi.ac.uk/interpro/) to search for protein functional domains. CBD, <u>c</u>yclic nucleotide-monophosphate <u>b</u>inding <u>d</u>omain (InterPro IPR000595); C2, C2 domain (InterPro IPR000008). Number under each domain indicate amino acids spanning these domains in TcCC2CP. B) Immunofluorescence microscopy of endogenously tagged *TcCC2CP*-3xc-Myc epimastigotes. From left to right: differential interference contrast (DIC), TcCC2CP-3xc-Myc (CC2CP, green), TcVP1 (VP1, red), Merge 1 image shows TcCC2CP-3xc-Myc (green) and TcVP1 staining (red), and Merge 2 shows TcCC2CP-3xc-Myc (green), TcVP1 staining (red) and DAPI labeling of nucleus and kinetoplast (blue). Co-localization of TcCC2CP and TcVP1 can be partially observed in merged images (yellow). Scale bar: 5 μm.

**Table S1.**
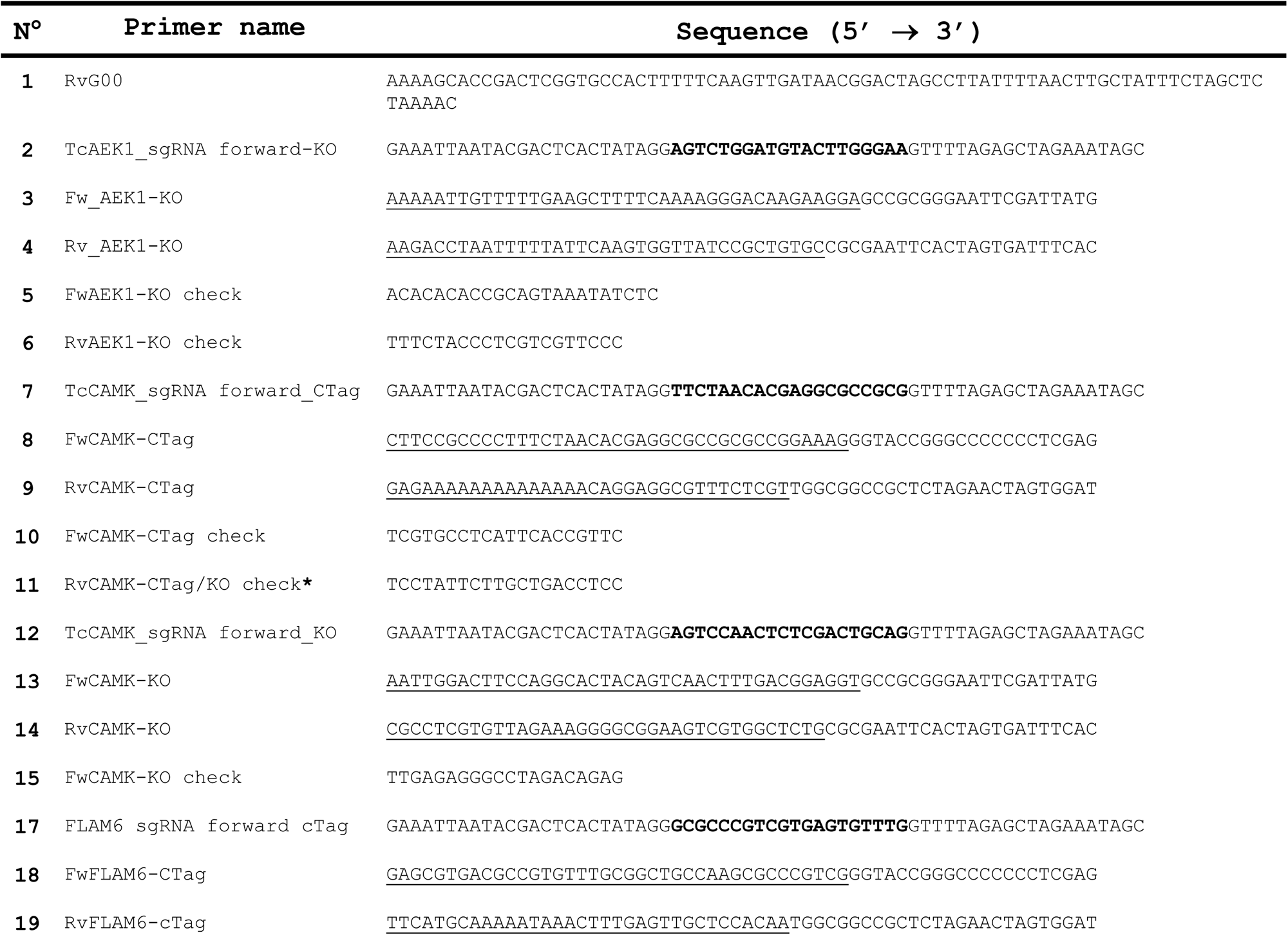
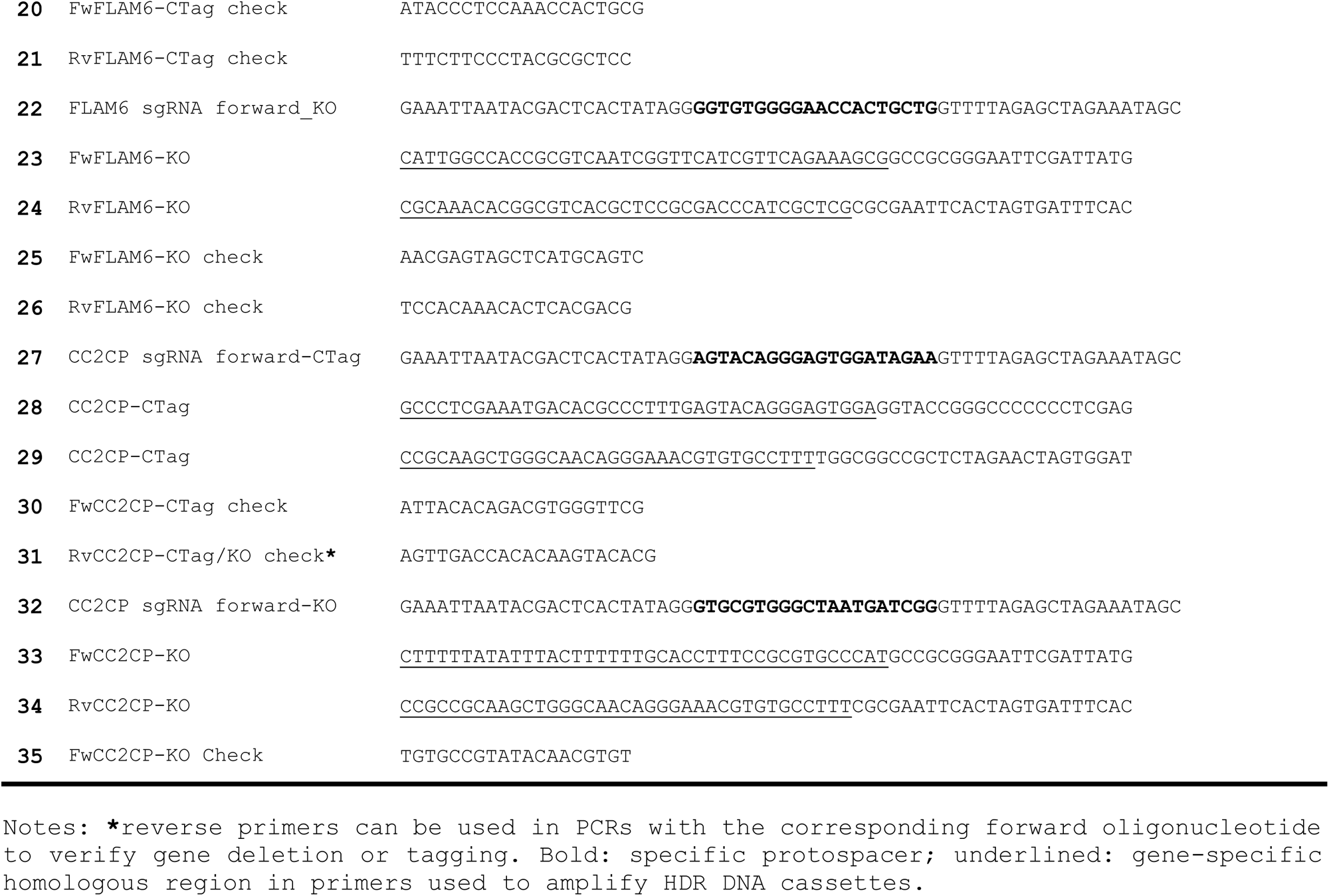
Oligonucleotides used in this work.

