## Supplementary figures and images for "Gene editing of putative cAMP and Ca^2+^-regulated proteins using an efficient cloning-free CRISPR/Cas9 system in *Trypanosoma cruzi*"

### Supplemental figures 1 and 2

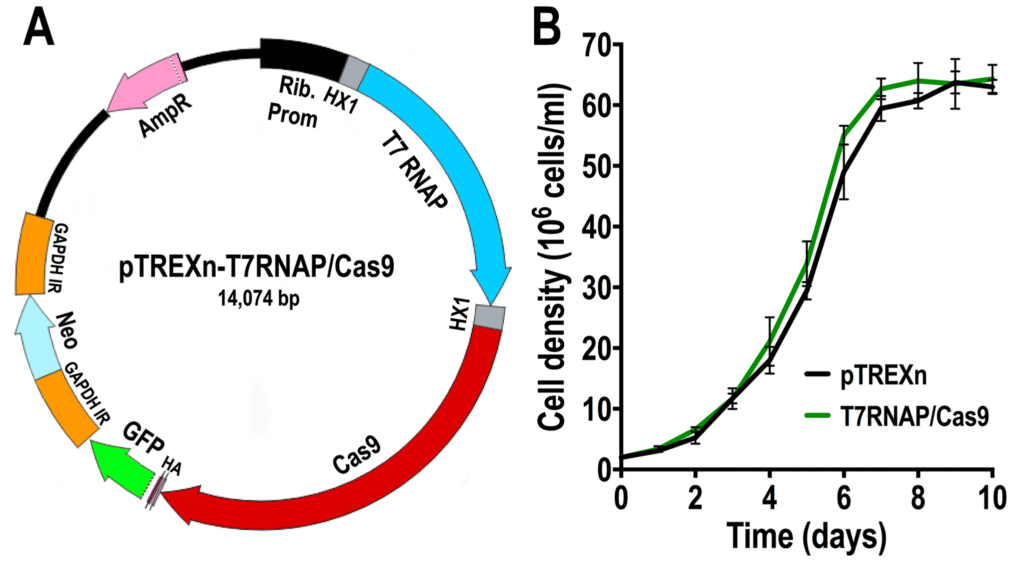

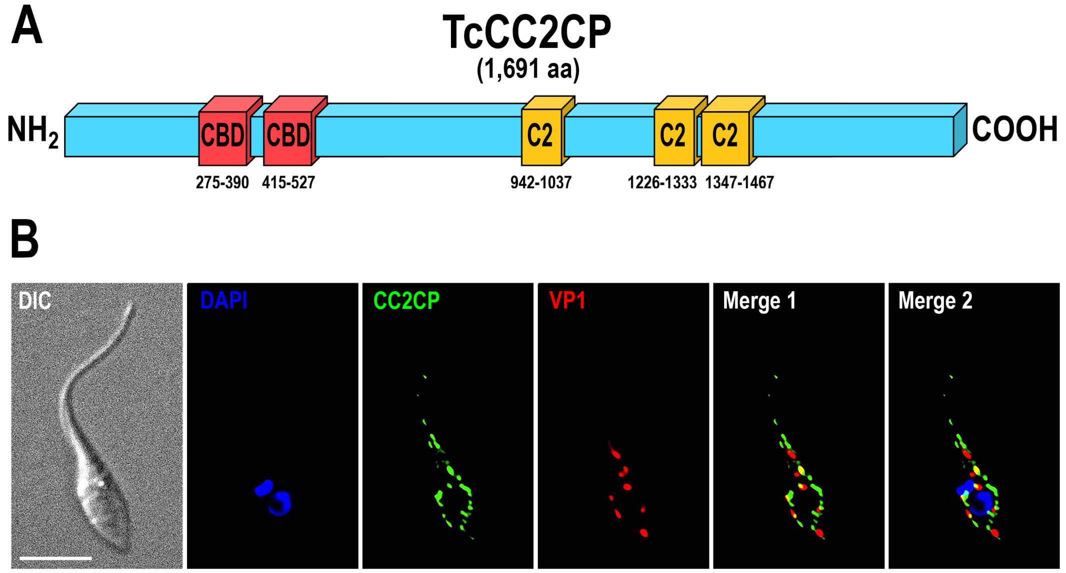
